# Estimating the rate of cell type degeneration from epigenetic sequencing of cell-free DNA

**DOI:** 10.1101/2020.01.15.907022

**Authors:** Christa Caggiano, Barbara Celona, Fleur Garton, Joel Mefford, Brian Black, Catherine Lomen-Hoerth, Andrew Dahl, Noah Zaitlen

## Abstract

Circulating cell-free DNA (cfDNA) in the bloodstream originates from dying cells and is a promising non-invasive biomarker for cell death. Here, we develop a method to accurately estimate the relative abundances of cell types contributing to cfDNA. We leverage the distinct DNA methylation profile of each cell type throughout the body. Decomposing the cfDNA mixture is difficult, as fragments from relevant cell types may only be present in a small amount. We propose an algorithm, CelFiE, that estimates cell type proportion from both whole genome cfDNA input and reference data. CelFiE accommodates low coverage data, does not rely on CpG site curation, and estimates contributions from multiple unknown cell types that are not available in reference data. In simulations we show that CelFiE can accurately estimate known and unknown cell type of origin of cfDNA mixtures in low coverage and noisy data. Simulations also demonstrate that we can effectively estimate cfDNA originating from rare cell types composing less than 0.01% of the total cfDNA. To validate CelFiE, we use a positive control: cfDNA extracted from pregnant and non-pregnant women. CelFiE estimates a large placenta component specifically in pregnant women (*p* = 9.1 × 10^−5^). Finally, we use CelFiE to decompose cfDNA from ALS patients and age matched controls. We find increased cfDNA concentrations in ALS patients (*p* = 3.0 × 10^−3^). Specifically, CelFiE estimates increased skeletal muscle component in the cfDNA of ALS patients (*p* = 2.6 × 10^−3^), which is consistent with muscle impairment characterizing ALS. Quantification of skeletal muscle death in ALS is novel, and overall suggests that CelFiE may be a useful tool for biomarker discovery and monitoring of disease progression.

## 1 Introduction

Cells die at different rates as a function of disease state, age, environmental exposure, and behavior [1][2]. Knowing the rate at which cells die is a fundamental scientific question, with direct translational applicability. A quantifiable indication of cell death could facilitate disease diagnosis and prognosis, prioritize patients for admission into clinical trials, and improve evaluations of treatment efficacy and disease progression [3][4][5][6]. Circulating cell-free DNA (cfDNA) is a promising candidate for understanding cell type specific death. When a cell dies, DNA is released into the bloodstream in short fragments (approximately 160bp, the length of a nucleosome) [7]. All people are thought to have a low level of cfDNA in their blood [8][9]. In healthy individuals, cfDNA in the blood likely arises from normal cell turnover. In individuals with a disease, cfDNA can come from illness specific apotosis and necrosis [10]. As a result, cfDNA levels have been shown to be elevated in individuals with cancer, autoimmune diseases, transplantation responses, and trauma [11][12][13][14]. CfDNA has become the clinical standard for non-invasive prenatal testing [15], and many companies and research groups are sequencing cfDNA to identify the presence of somatic mutations related to tumors [16][17][18].

To understand what drives changes in the biology of people with disease, we can decompose the cfDNA mixture into the cell types from which the cfDNA originates. This can give a non-invasive picture of cell death, which can be used to characterize an individual’s disease, or health, at a particular moment. While each cell type has the same DNA sequence, which does not give us information on where a cfDNA fragment arises from, DNA methylation is cell type specific [19]. Subsequently, there is a rich literature of cell type decomposition approaches using DNA methylation, often focusing on estimating the contribution of immunological cell types to whole blood [20][21][22][23].

More recent work has attempted to use cfDNA methylation patterns to decompose tissue of origin for cfDNA [24][25][26][27]. These approaches, however, do not address some of the unique challenges of cfDNA. Previous work was designed for reference and input data from a methylation chip, which are high coverage and have relatively low noise. Since cfDNA is only present in the blood in small amounts, as an onerous amount of blood must be extracted from a patient to get the required amount of input DNA for methylation chips, which may not be practical for clinical use [28]. Other technologies and methods focus on sensitive detection of specific tissues or cancer sites [29][30][31]. While increasingly powerful, these approaches can not provide biomarker discovery or comprehensive deconvolution. In this work, we turn to using whole genome bisulfite sequencing (WGBS) to assess the methylation of cfDNA. Unlike methylation chips that target specific genomic locations, WGBS covers the entire genome, typically resulting in lower coverage per-site, and increased noise relative to chip data. Current methods are ill-equipped to handle such noise in either the reference or input.

Previous methods are also limited by which DNA methylation sites (CpGs) are chosen. Methylation chips survey a limited number of CpGs, which may not be maximally informative of cell types. Some approaches also rely on selecting a set of CpGs designed for a particular dataset [24][26]. While curated site selection is useful for specific biological queries, it can cause bias when generalizing to other settings or diseases. Choosing which sites to include in a decomposition can substantially influence which cell types are predicted, because different sites are informative for different cell types. Another important limitation of previous cfDNA decomposition methods is that the results are restricted to the cell types included in the reference panel. The decomposition using these methods estimates the cfDNA mixture as being exactly composed of the cell types given in the input. However, as there are many hundreds of cell types throughout the body, it is currently impossible to incorporate all or most into a reference panel. Thus, the specific choice of reference cell types will lead to obvious biases in decomposition results of these methods.

In this work, we develop an efficient EM algorithm, CelFiE (CELl Free dna Estimation via expectation-maximization) for cfDNA decomposition that allows for low coverage and noisy data. Our method can also estimate unknown cell types not included in a reference panel and is not dependent on curated input methylation sites. We show in realistic simulations that we can accurately estimate known and unknown cell types even at low coverage and relatively few sites. We also can estimate rare cell types that contribute to only a small fraction of the total cfDNA. Decomposition of real WGBS complex mixtures demonstrates that CelFiE is robust to several violations of our model assumptions. Specifically, the real data allow: correlation across regions and between cell types, read counts that are drawn from non-uniform distributions, and reference samples that are actually heterogeneous mixtures of many cell types. Additionally, we develop an approach for unbiased site selection.

Finally, we apply CelFiE to two real cfDNA data sets. First, we apply CelFiE to cfDNA extracted from pregnant women and non-pregnant women. Since placenta is not expected in non-pregnant women, this data enables validation for our method. We also apply CelFiE to cfDNA from amyotrophic lateral sclerosis (ALS) patients and age matched controls. Currently, there are no established biomarkers for ALS. Subsequently, it is difficult to monitor disease progression and efficiently evalute treatment response [32]. CfDNA provides an opportunity to measure cell death in ALS that could fill these gaps. We find a significantly elevated skeletal muscle component in ALS patients. This novel observation, along with successful decomposition of cfDNA from pregnant women, demonstrates that CelFiE has the potential to meaningfully decompose cfDNA in realistic conditions. CfDNA decomposition has broad translational utility for understanding the biology of cell death, and in applications such as quantitative biomarker discovery or in the non-invasive monitoring and diagnosis of disease.

## 2 Methods

The objective of this work is to estimate the relative proportions of various cell types that contribute to the cfDNA of an individual. We assume that we are provided with a bisulfite sequenced reference data set, comprised of *T* cell types indexed by *t*, at *M* CpG sites indexed by *m*. Bisulfite sequencing produces read counts from specific cell types that we collect in two *T* × *M* matrices: *Y* and *D*^*Y*^, where, *Y*_*tm*_ and 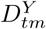 are the number of methylated and total reads at CpG *m*, respectively, in reference cell type *t*. Together, these two matrices represent our reference cfDNA data.

We are also provided with cfDNA extracted from *N* individuals indexed by *n*. The bisulfite sequencing read counts of the cfDNA are given in two *N* × *M* matrices *X* and *D*^*X*^, with *X*_*nm*_ and 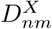 giving the number of methylated and total reads at CpG *m* in the cfDNA from individual *n*, respectively. These two matrices represent our sample cfDNA data.

Our algorithm, CelFiE, takes as input the matrices *Y*, *D*^*Y*^, *X*_*nm*_, and 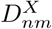, and then outputs a matrix *α*, where *α*_*nt*_ is the fraction of the cfDNA in person *n* that originated in cell type *t*. CelFiE also learns cell type-specific methylation proportions, which can improve on the reference proportions when they are noisy. Our algorithm permits *t* > *T*, meaning that we can estimate cell types that are not provided in the reference panel. This is mathematically equivalent to appending all-zero rows to *Y* and *D*^*Y*^, i.e. reference cell types with zero total coverage.

### 2.1 Model

We model the cfDNA as a mixture of DNA from cell types in the reference panel and, potentially, unknown cell types absent from the reference panel. We assume that the individuals are independent given the true, unknown methylation proportions of each cell type, and the individual-specific cell type proportions.

We assume that reference data are drawn from a binomial distribution:

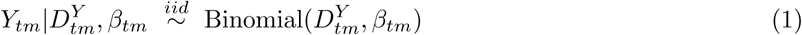

where *β*_*tm*_ ∈ [0, 1] is the true, unknown proportion of DNA in a cell type that is methylated at position *m*. This model assumes no intra-cell type heterogeneity, in the sense that each cell in a cell type has identical methylation probability.

Next, we model the samples in the cfDNA data. We assume each cfDNA read is drawn from some cell type *t* at some marker *m*, and in turn that its methylation value is drawn from a Bernoulli distribution governed solely by the methylation proportion in the cell type of origin:

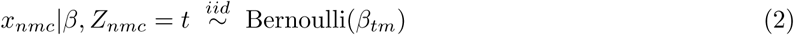

where *x*_*nmc*_ is the methylation status of the *c*-th read from sample *n* at position *m*, and *Z*_*nmc*_ = *t* indicates that *t* is the cell type of origin for this read.

For each person and methylation site, we define the total number of methylated reads as 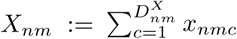. This simply sums the methylation status over all reads for each person at each site. In the special case where 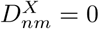, we define *X*_*nm*_ = 0.

Finally, we assume that the cell type of origin of each cfDNA molecule is drawn independently from some individual-specific multinomial distribution:

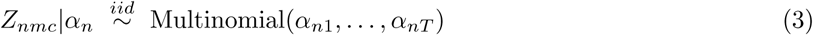

where *α*_*nt*_ is the probability that a read from person *n* comes from cell type *t*.

### 2.2 EM algorithm for one cfDNA sample

For simplicity, we first describe CelFiE in the case where the cfDNA data set contains only a single person, meaning the decomposition relies almost exclusively on the reference panel. We then explain how CelFiE can jointly model multiple individuals in the cfDNA data, as well as how and why this enables the estimation of unknown cell types. Full details of both algorithm derivations are given in the Appendix.

Formally, assume there is only one sample in the cfDNA data (i.e. *N* = 1). We define *z*_*tmc*_ as a binary indicator for whether read *c* at CpG *m* for the single cfDNA individual originates from cell type *t*. In relation to *Z* above, *z*_*tmc*_ = 1 if *Z*_1*mc*_ = *t*, and otherwise 0. That is, *Z*_1*mc*_ is a categorical variable, and *z*_*tmc*_ indicates which value *Z*_1*mc*_ takes.

To calculate the full data likelihood, *P* (*x, z, Y* |*α, β*), we first factorize it into *P* (*x, Y* |*z, α, β*) · *P* (*z* |*α, β*). This then simplifies into three components:

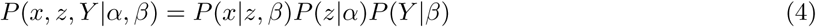

The first component defines the probability of the cfDNA reads, given which cell type they come from and the methylation proportions of those cell types. The third component analogously defines the probability of drawing the reference reads. The second component describes the probability of observing a specific cell type in the cfDNA, which is determined by the proportion of each cell type in the individual’s cfDNA. We show in the Appendix that the resulting log-likelihood is equivalent to:

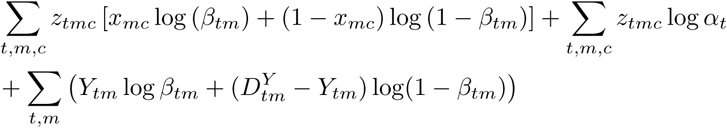

For this one-sample section, we drop an index on *x* and write *x*_*mc*_ instead of *x*_1*mc*_. Analogously, we write *X*_*nm*_ = *X*_*m*_ as the total number of methylated reads at position *m* (and 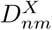 as 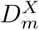).

To calculate the expected log likelihood, i.e. the *Q* function, we must integrate over the conditional distribution for the missing data, i.e. *P* (*z*|*x, β, α*). Since *z*_*tmc*_ is binary and each read and site is assumed independent, this distribution is the probability that each *z*_*tmc*_ is 1. In other words, the probabilities that each read comes from each cell type are sufficient statistics, and are given by (Section 7):

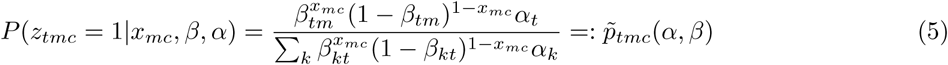

Conceptually, if read *c* is methylated, this indicates the read is more likely to come from cell types with high methylation proportion, as *β*_*tm*_ is larger (and vice versa if the read is unmethylated). Regardless the methylation state, however, this equation also says that the read is likelier to come from more common cell types, as *α*_*t*_ is larger.

This final term 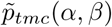, seems complex. However, it actually only depends on the specific read *c* through its methylation status, and takes only two values. We can redefine it in simpler terms, which represents the probability of each cell type for each read depending on its methylation:

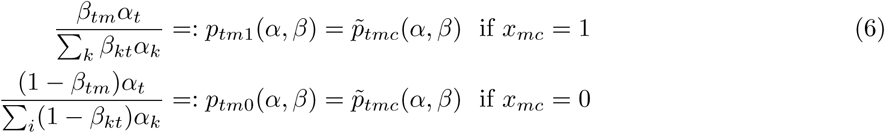

**E step:** The *Q* function is defined at iteration *i* by:

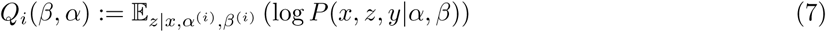

where *α*^(*i*)^ and *β*^(*i*)^ are the parameter estimates of the cell type proportions and methylation proportions from the last EM step. Let 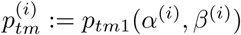, which is the probability that a methylated read at site *m* comes from cell type *t* given the previously estimated parameters from iteration *i*. Then *Q*_*i*_ is (Section 7):

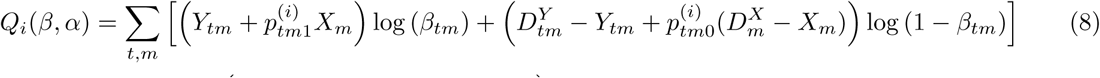

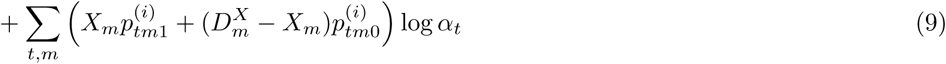

The first line in this equation captures the expected total number of methylated reads (first term in the sum) and the total number of expected unmethylated reads (second term) for each cell type and site. Each of these terms combines both the reference and cfDNA contribution, e.g. the first term combines the total methylated reads from the relevant reference cell type (*Y*_*tm*_) with the expected number of methylated reads from that cell type in the cfDNA mixture 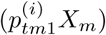.

Complementary to the first line, the second line determines the likelihood of *α* and does not depend on *β*. It captures the likelihood of observing the expected cell type frequencies. This is given by the sum of the expected methylated and the expected unmethylated reads over all loci.

**M step:** To update the estimated cell type proportions, *α*, we maximize *Q*_*i*_ under the constraint that *α* is a probability vector, i.e. its entries are non-negative and sum to one. The maximizer is (Section 7):

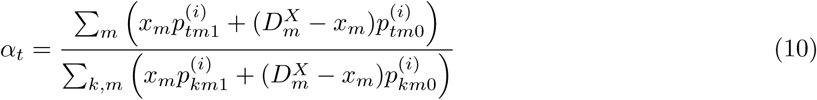

The numerator is simply the number of reads expected to originate from each cell type, which is calculated by adding the expected contributions from the methylated and the unmethylated reads. The proportions are then obtained by normalizing these numerators to sum to 1.

The other M step update is for *β*, the proportion of reads that are methylated at each site and in each cell type (Appendix):

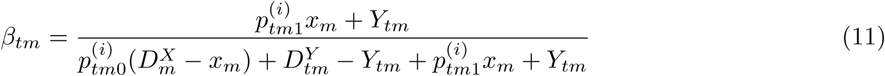

Intuitively, this is the ratio of the expected number of methylated vs total reads from cell type *t* at site *m*. This update is conceptually similar to the *α* update in the sense that it matches an estimated proportion to an expected proportion. For *α*_*t*_, this is the expected proportion of reads deriving from cell type *t*; for *β*_*tm*_, this is the expected proportion of reads from cell type *t* that are methylated at site *m*.

### 2.3 EM algorithm for multiple cfDNA samples

We now return to allowing *N* > 1 cfDNA samples. In this setting, *α* is a matrix, because each cfDNA sample may have different proportions of each cell type in their cfDNA mixture. Further, *x*_*nmc*_ and *Z*_*nmc*_ are now 3-dimensional arrays indexed by cfDNA individual *n*, methylation site *m*, and sequencing read *c*, and the binary indicators *z*_*nmtc*_ are now 4-dimensional, as they additionally index each cell type.

The conditional distribution for *z* at each step of the EM algorithm now becomes:

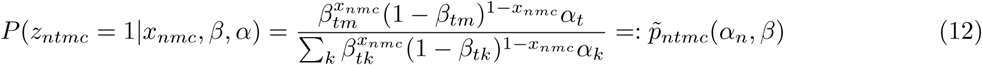

As before, this 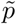 term depends on *c* only through *x*_*nmc*_, and so we simplify terms by defining 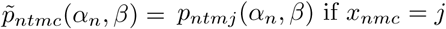 if *x*_*nmc*_ = *j* for *j* = 0, 1.

To simplify the E step, we define the responsibilities by 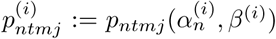. For *j* = 0, this gives the conditional probability that an unmethylated read from individual *n* as site *m* comes from cell type *t* given the current parameter estimates; *j* = 1 gives the analogous probability for methylated reads. Since we assume cfDNA individuals are independent given *α* and *β*, the E step is a simple generalization of the one-sample E step that sums over samples and can be written:

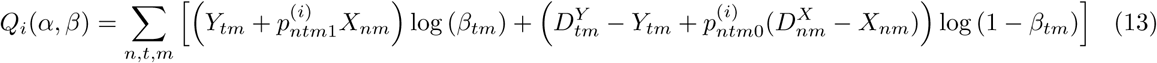

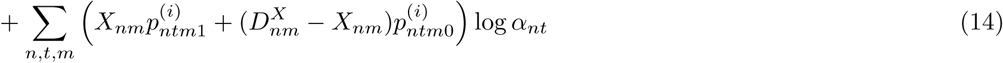

This *Q* function can be interpreted identically to the single-sample *Q* function. The only difference is that now reference reads are added with expected cfDNA reads for multiple individuals, and the expectations 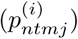 depend on cfDNA individual *n* as well as cell type *t*, CpG site *m*, and methylation status *j*.

*Q*_*i*_ additively splits over row of *α*, therefore, the updates for each *α*_*n*,_ are identical to the single-sample *α* updates, where *α*_*nt*_ replaces *α*_*t*_, *X*_*nm*_ replaces *X*_*m*_, 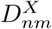 replaces 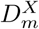, and 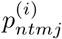 replaces 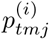. This means that if we condition on the number of reads coming from each cell type in person *n*, our estimates of that person’s cell type proportion does not depend on anything else.

For *β*_*tm*_, the M-step again compares the expected number of methylated and unmethylated reads at CpG *m* from cell type *t*, where the expectation combines reads from reference cell type *t* with the expected number of cfDNA reads from cell type *t*. The only difference is that now the expectation combines the expected contributions from multiple cfDNA samples:

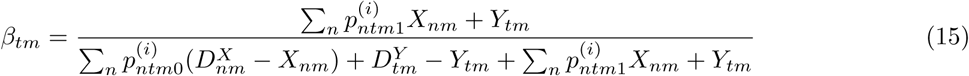

#### Unknown sources

It is likely that there are cell types in the cfDNA mixture not contained in the reference data. To estimate the proportion of an unknown cell type with CelFiE, we append a zero row to *D*^*Y*^ and *Y*, and then run CelFiE as usual. This produces an EM that is mathematically similar to the STRUCTURE model of mixtures of human populations [33]. Essentially, CelFiE estimates methylation patterns and abundances for the unknown cell type(s) that maximize the overall likelihood. To model more than one unknown cell types, additional rows of zeros are added to *D*^*Y*^ and *Y*. Note that if the number of unknown cell types is greater than the number of individuals, the problem is not identified.

#### Regularization and Missing Data

Missing observations are allowed in both the reference and the input. It is represented as a 0 entered in both *X*/*D*^*X*^ or *Y* /*D*^*Y*^. In practice, we add a methylated and unmethylated pseudocount to every entry of *X* and *Y* /*D*^*X*^ and *D*^*Y*^ to stabilize the algorithm and likelihood in case of cell type/site combinations with very low coverage.

#### Computational cost

Each iteration of the EM algorithm in CelFiE involves three calculations. First, 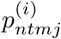 is evaluated for each sample *n*, cell type *t*, CpG site *m*, and methylation status *j* = 0, 1; each calculation is independent of the input data dimensions, hence evaluating *p*^(*i*)^ is *O*(*NTM*). Second, *α*_*nt*_ must be evaluated, which involves summing over *M* sites for each *n* and *t*, giving overall complexity *O*(*NTM*). Finally, updating *β*_*tm*_ requires summing over all cfDNA individuals and the reference cell type data, again giving overall complexity *O*(*NTM*). Overall, this means that CelFiE scales linearly in sample size, number of CpGs, and number of cell types.

We also note that if multiple references were included, the cost would not multiply–rather, the cost would increse to *O*((*N* + *N*_*ref*_) *TM*), where *N*_*ref*_ is the (maximum) number of reference samples per cell type.

## 3 Results

### 3.1 Evaluation using simulated cfDNA mixtures

We began by simulating cfDNA mixtures informed by realistic sequencing conditions: namely, low read count and noisy. The results of CelFiE in these conditions are compared with MethAtlas for context [25]. While MethAtlas is not designed for low read count data, it is, to the best of our knowledge, the only other cfDNA decomposition algorithm that allows inclusion of arbitrary input sites, and does not restrict to specific cell types in the reference. To match the reference data provided in MethAtlas, we simulated 8,000 CpGs and 25 cell types [34]. The true methylation proportions of each CpG was drawn independently from a uniform distribution, so that the methylation of each CpG was between 0% and 100%. To characterize the decomposition performance of CelFiE across both rare and abundant cell types, we defined the true cell type proportion vector as 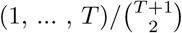, where *T* = 25 is the number of cell types truly in the mixture. For CelFiE, the input data is the number of methylated reads and read depth at each site. The reference read depths were drawn independently from a Poisson distribution centered at 10, a relatively low sequencing depth for a WGBS experiment [35]. The number of methylated reads for a given CpG in each of the 25 cell types was drawn from a binomial distribution, where the probability of success was the true methylation value in that cell type, and the number of trials was the read depth at that locus. CfDNA read depths for each CpG were simulated from a Poisson distribution centered at 10, and then the reads for each CpG were assigned to originate from a cell type based on the cell type proportion vector for the cfDNA mixture. A read was determined to be either methylated or unmethylated given the true methylation proportion in that read’s cell type of origin at that CpG. Since MethAtlas was designed for chip data, for the MethAtlas reference data, we calculate the methylation proportion for a CpG by dividing the methylated reads by the depth at that locus.

In total, we perform 50 independent simulations of both CelFiE and MethAtlas. For each of these simulations, we generate data as described above. We perform 10 random restarts of the EM algorithm per simulation to avoid getting stuck in local optima. We choose the best random random restart by choosing the restart with the largest log likelihood.

As expected, in this experiment CelFiE performs better than MethAtlas at low read depths (Figure 1). Per simulation, we calculate the Pearson’s correlation between the true cell type proportion vector and the estimated proportions vector. For MethAtlas the mean *r*^2^ across replicates is 0.22, while CelFiE’s mean *r*^2^ is 0.96.

**Fig. 1:**
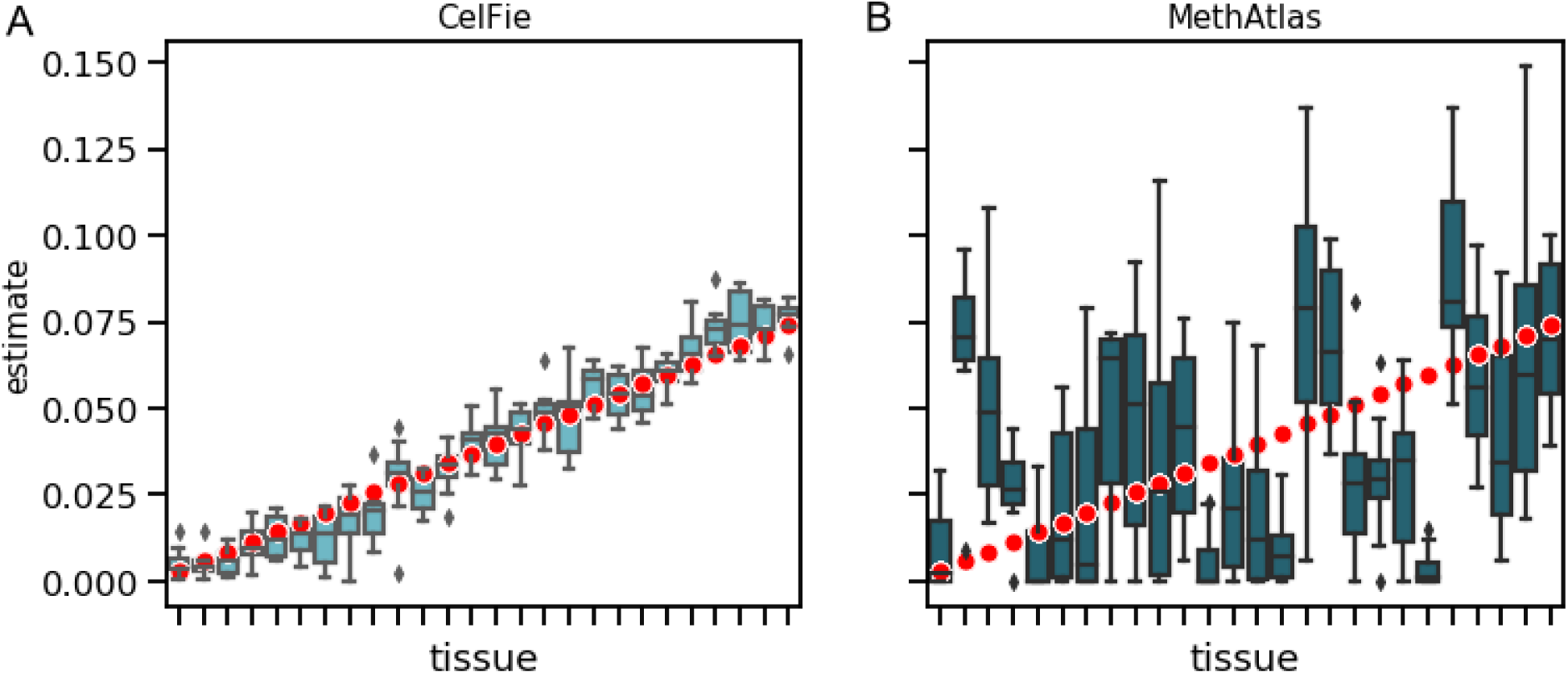
Decomposition of simulated cfDNA mixtures by CelFiE (A) and MethAtlas (B). 50 replications were and the estimated mixing proportions were plotted (light blue and dark blue boxes, respectively). The true cell type proportions are depicted as red points.

After demonstrating CelFiE’s performance in context with MethAtlas, we further characterized the properties of our approach. We make two modifications to the basic simulation in Figure 1. First, we vary the number of CpGs (100, 1,000 and 10,000), which represents conditions with varying amounts of information about cell type provided to our algorithm. Second, we focus on a single cell type and vary its proportion between 0% and 100%. In total we simulate 10 tissues, and fix one tissue. The remaining 9 additional cell type proportions are drawn from an independent uniform distribution and then renormalized so all 10 proportions sum to one.

Performance is assessed by calculating the Pearson’s correlation between the estimated cell type proportions and the true proportions for 50 replicates. We find that as the number of sites increases, the capability of CelFiE to accurately decompose the cfDNA mixtures improves (Figure 2A), especially at less abundant cell types.

**Fig. 2:**
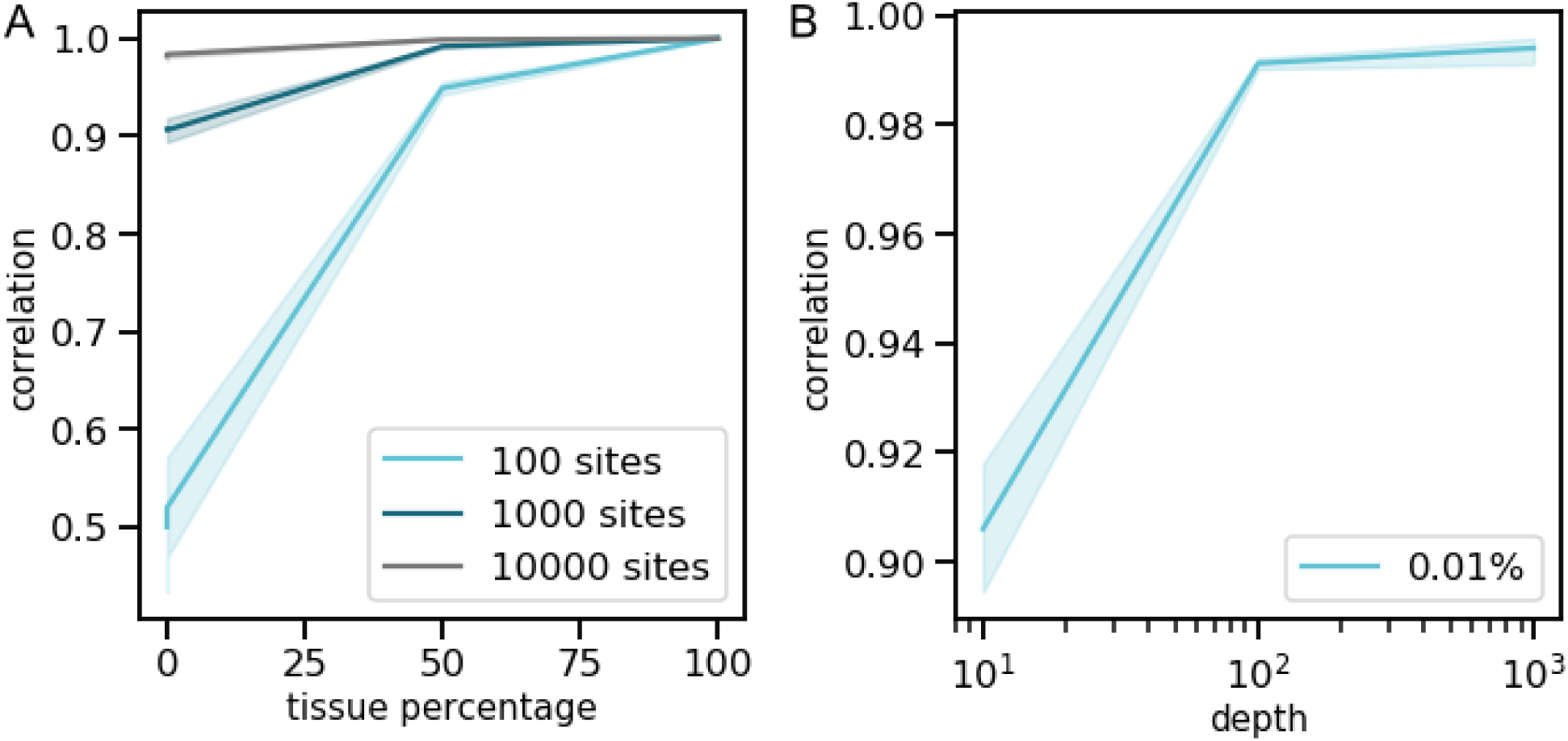
Pearson’s correlation between real and estimated cell type proportion. We fix a cell type at a proportion between 0% and 100%, and estimate the cell type proportion for 100, 1000, and 10000 sites (A). For a rare cell type of 0.01%, we plot the Pearson’s correlation between the true and estimated proportion using 1000 CpG sites at a range of depths (B).

Previous work suggests that a large portion of cfDNA originates from white blood cells [24]. This suggests that a potential cell type of clinical significance may only be present at a low proportion in the mixture. To assess the ability of CelFiE to estimate rare cell types, we simulate data as in Figure 2A, but fix one cell type at 0.01%, a very low abudance. In order to understand how CelFiE’s ability to estimate rare cell types changes as a function of sequencing depth, we simulated input and reference reads at 10x, 100x, and 1000x coverage for 1000 CpGs. We then calculate the correlation between the estimated cell type proportions and the true proportions. Even at this low read depth, CelFiE can decompose extremely low incidence cell types (Figure 2B).

Next, we turned to understanding the behavior of CelFiE when estimating unknown cell types. We continue simulating data with low read counts: creating reference and cfDNA reads for 1000 CpGs at 10x depth as in Figure 1. For the cfDNA mixtures, ten cell types were truly in the mixture. One cell type was excluded from the reference and fixed at 20% of the cfDNA mixture, an abundant proportion. We simulate cfDNA reads for 1, 5, 10, 50, 100, and 500 individuals, as the unknown component will over-fit when the number of individuals is too low to properly learn the missing tissue. For these simulations, we assume that each individual has the same cell type proportions. We then calculate the average relative mean squared error (MSE) for the unknown component, defined as the MSE between the true unknown proportion and the average estimated proportion across all individuals, divided by the true proportion. As the number of people included in the decomposition increases, the performance of CelFiE improves (Figure 3A).

**Fig. 3:**
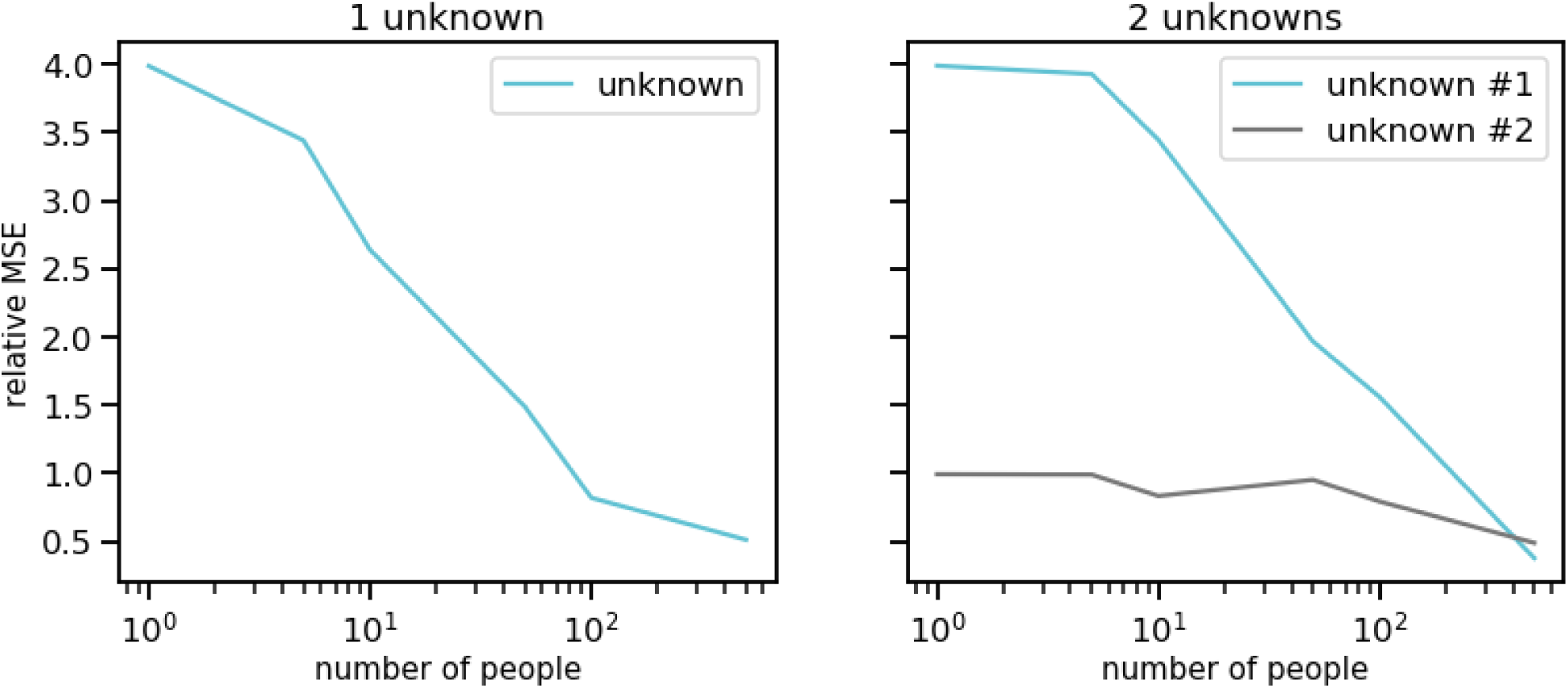
Decomposition results for cfDNA mixtures with missing cell types in the reference. We simulate cfDNA for 1, 5, 10, 50, 100 and 500 people, and exclude one cell type truly in the mixture at 20% (light blue) (A) or two cell types (B), one in the mixture at 20% (light blue), and the other at 10% (dark blue). We calculate the relative mean square error, defined as the true unknown proportion and the average estimated proportion across all individuals, divided by the true proportion for 50 simulation experiments.

We then exclude two abundant cell types, fixing one at 20%, and one at 10%. Since the inferred CelFiE labels are not identified (i.e. CelFiE’s estimated unknown proportion 1 can correspond to either missing cell type 1 or 2), we assigned the larger estimated component to correspond to the 20% missing cell type, and the smaller to the 10% missing cell type. We then calculate the relative MSE between the true unknown proportion and the estimated proportion. When two cell types are excluded from the reference, more people are needed to accurately estimate the 20% unknown component (Figure 3B).

Our ability to accurately estimate unknowns contrasts with MethAtlas, which can only estimate proportions for cell types in the reference. This creates bias in the decomposition that can be addressed with CelFiE. Specifically, if we simulate cfDNA mixtures as in Figure 1, but one cell type is excluded from the reference, MethAtlas overestimates the cell type proportions (see Figure S3). Nonetheless, as expected, MethAtlas and CelFiE perform similarly at high read depths and when all cell types are known (Figure S4). Overall, CelFiE is a desirable alternative to MethAtlas for low sequencing depths and an incomplete reference panel.

### 3.2 Performance on WGBS cfDNA mixtures

After demonstrating the performance of CelFiE in pure simulations with realistic parameters, we consider mixtures made from real WGBS data. These data are substantially more complex and violate several assumptions of the CelFiE algorithm. In particular, the reference data contain tissues composed of multiple cell types, CpGs are correlated locally across genomic regions and between cell types, and read counts are drawn from non-uniform distributions that reflect true biological and technical heterogeneity across sites. To examine how robust CelFiE is to these complications, we use biological replicates for 10 WGBS data sets (small intestine, pancreas, monocyte, stomach, tibial nerve, macrophage, memory B cell, adipose, neutrophil, and CD4 T cell), downloaded from the ENCODE and BLUEPRINT projects [36][37][38]. These data encompass both and cell types and tissues, which are essentiallly complex mixtures of cell types. In all experiments, we chose to include tissues to see if their differential cell type mixtures contribute to error in our decomposition results. One set of WGBS biological replicates was assigned to make up the cfDNA mixtures; the other was assigned to the reference matrix. We then generated cfDNA mixtures for 100 individuals.

Since roughly 80% of CpG sites in the human genome do not vary between cell types [39], randomly selected CpGs will be contain mostly uninformative loci. To limit uninformative CpGs included in our analysis, we developed a method for choosing a set of unbiased informative CpGs, which we call tissue informative markers (TIMs) (Section 7.5). We selected 1,060 TIMs, or approximately 100 TIMs per WGBS sample, for use in these simulations. Selecting TIMs improves performance over a random subset of CpGs (Figure S1). Furthermore, DNA methylation of nearby CpGs is correlated [40]. We thus combine related CpG sites to increase read depth and overcome noise (Section 7.5).

After TIM selection, we subset the WGBS samples to these TIMs, and we combine reads for all CpGs 250bp upstream and downstream of the TIM. We create complex mixtures of the WGBS samples according to 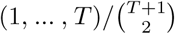, where *T* = 10. For these complex mixtures of cfDNA at our selected TIMs, CelFiE estimates cell type methylation values with an average Pearson’s *r*^2^ = 0.97 (Figure S5A) to the true WGBS data. The high correlation demonstrates potential value for downstream analyses, such as biomarker discovery. We can also successfully decompose these complex WGBS mixtures into their cell type proportions (Figure 4A).

**Fig. 4:**
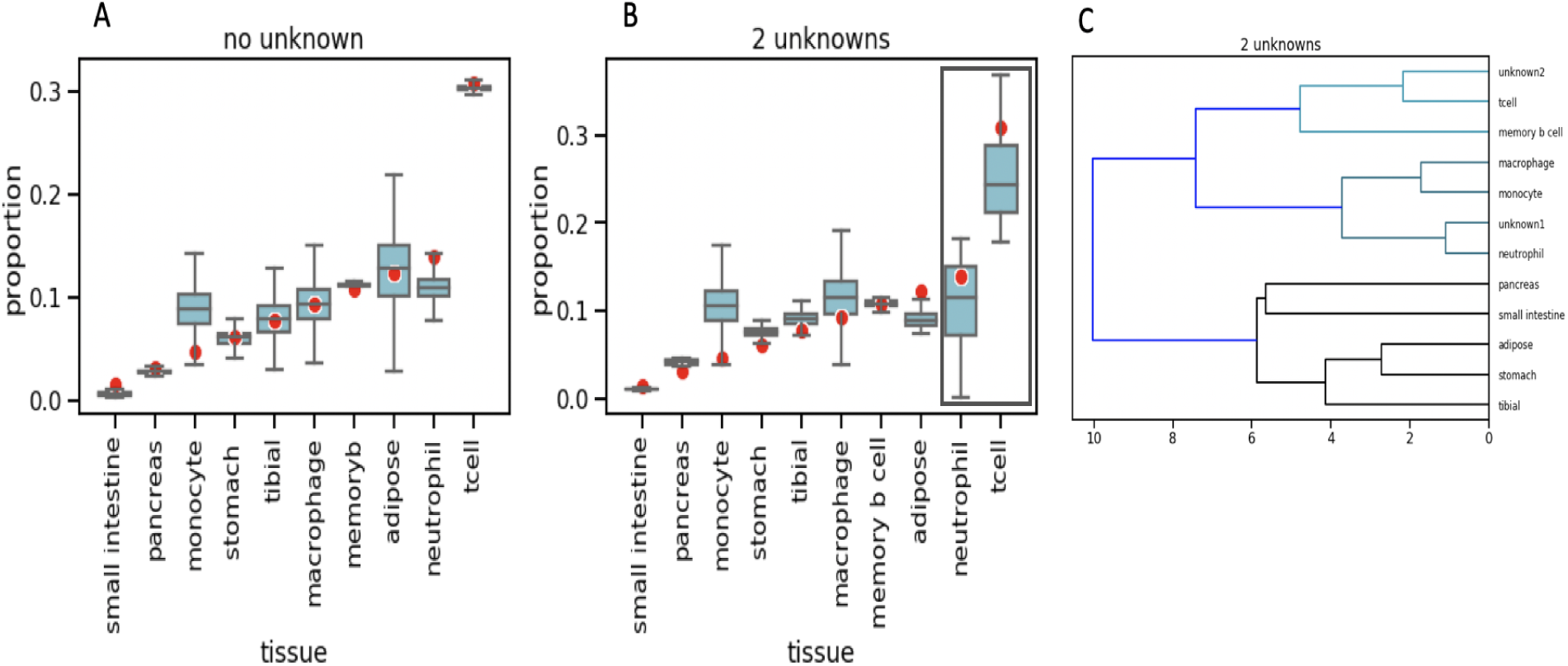
CelFiE cell type proportion estimates for real WGBS cfDNA mixtures. We estimate (blue boxes) the true cell type composition (red dots) for 100 individuals in 50 simulation experiments in the scenario where there are no missing cell types (A) and when two cell types are missing from the reference (indicated by the grey box) (B) and (C) hierarchical clustering of the estimated methylation proportions for the two missing cell types, where unknown 1 was neutrophil and unknown 2 was t-cell.

We next consider the more realistic scenario of missing cell types. To investigate our ability to estimate mixtures with a substantial unknown component, we mask two cell types, neutrophil and T-cell, that are truly in the mixture from reference panel. T-cell is at a high proportion of 30%, and neutrophils are at a lower proportion, 14%. Using the summed TIM sites, we perform simulations for 100 people with identical cell type proprotions. We find that our ability to successfully decompose a complex mixture decreases when there are two missing cell types (Figure 4B). The estimated correlation to the true WGBS methylation values, however, remains high, with an average Pearson’s *r*^2^ = 0.93 (Figure S5B).

To further validate CelFiE’s ability to estimate missing cell types, we assessed how similar the learned methylation proportions for the missing cell types are to the true WGBS proportions for neutrophil and t-cell. To do this, we appended the methylation patterns learned by CelFiE of the two unknown cell types to the matrix of true reference methylation values, including the values for neutrophil and t-cell that were originally masked. We calculated a distance matrix for the reference matrix plus unknowns, and used this to perform hierarchical clustering. Figure 4C shows that the unknown cell types cluster with their true cell type. Specifically, unknown 1 clusters with the reference neutrophil sample, amongst other myeloid cell types, and unknown 2 clusters with the reference t-cell sample. Together with Figure 4B, this suggests that even with an incomplete reference, CelFiE estimates both the correct cell type proportion and cell type methylation values.

### 3.3 Application to real data

### 3.4 Application to real pregnancy data

Finally, we applied CelFiE to real cfDNA data. To validate our model, we choose pregnancy data because it is an example of a patient population with a reliable true positive [41]. Unlike decomposition of cell types in blood, there is no FACS or similar existing gold standard. Nonetheless, we know *a priori* that non-pregnant women will not have placenta cfDNA in their bloodstream.

To do this, we download publicly available WGBS cfDNA of 7 pregnant and 8 non-pregnant women [42]. All women were between 11 and 25 weeks pregnant at time of cfDNA extraction. We subset the WGBS sites to the same TIMs we use in Section 3.2 and sum all reads +/- 250 bp around each TIM (Section 7.5). Twenty WGBS datasets from the ENCODE and BLUEPRINT projects were chosen for the reference panel, representing tissues and cell types throughout the body and blood, along with one unknown category. We choose the random restart with the highest log likelihood of 10.

CelFiE estimates a high proportion of white blood cells, consistent with previous estimates based on cfDNA [25]. We use a single unknown cell type component, and we estimate that it is large (Figure 5). This suggests there is substantial cfDNA signal that cannot be captured by existing methods.

**Fig. 5:**
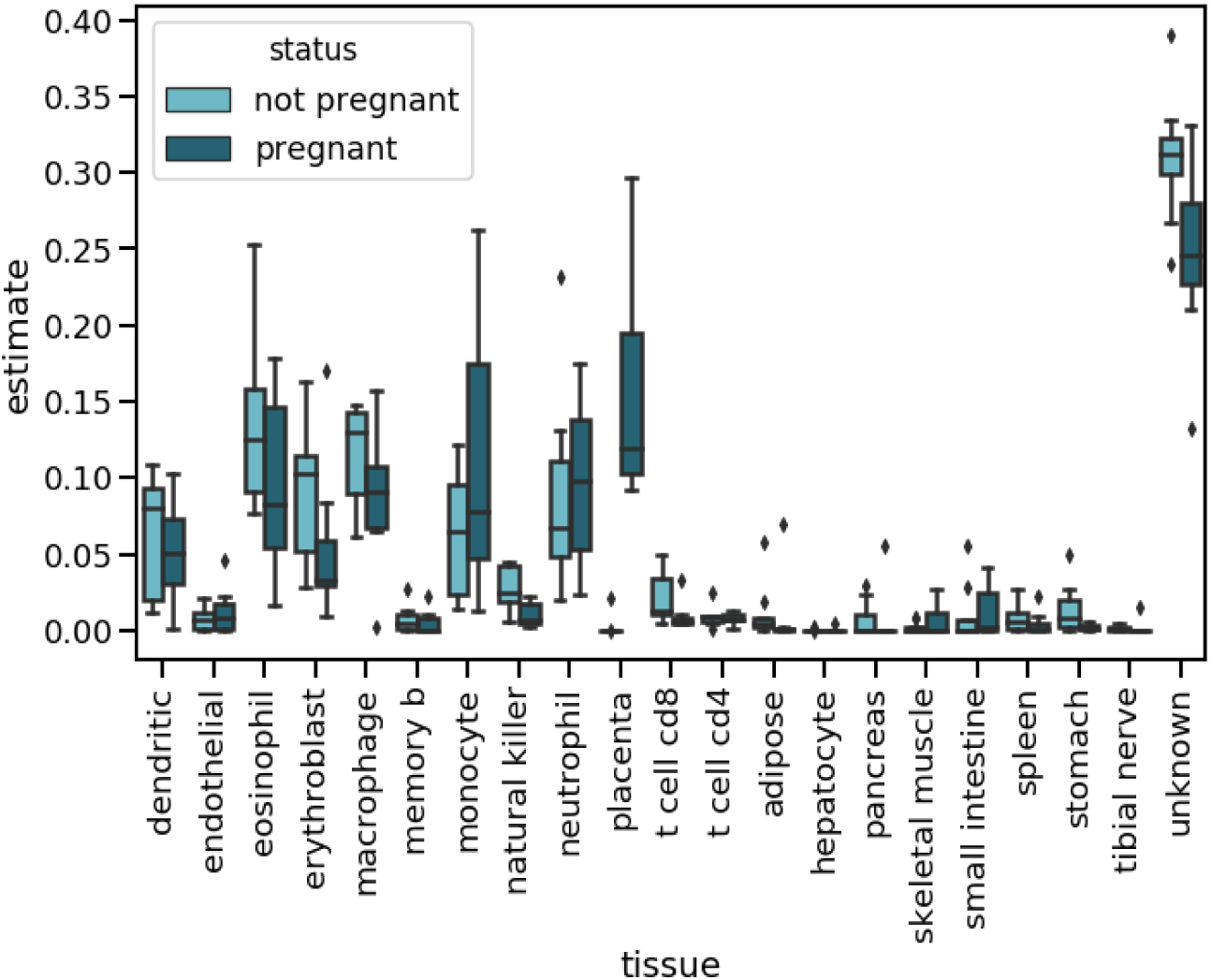
CelFiE decomposition estimates for 8 non-pregnant (light blue) and 7 pregnant women (dark blue).

To evaluate which cell types differ most between pregnancy states, we perform grouped two-sample *t*-tests of inferred cell type proportions. As expected, placenta shows by far the greatest difference, ranging from 9.3% to 29.7% (median 11.9%), and 2.9 × 10^−16^ to 2.1 × 10^−2^ (median 2.3 × 10^−12^) in pregnant and non-pregnant women, respectively (grouped t-test, *β* = 4.55, *s.e* = 0.81, *p* = 9.1 × 10^−5^). This is consistent with previous estimates of the proportion of placenta in the cfDNA (median 15.3% in trimester 1 and 2) [43]. We restrict statistical tests to the relevant tissue, in this case placenta, however, estimates are provided for all tissues and cell types in Figure 5. An important caveat for these tests is that they treat the inferred cell type proportions as though they are known, which fails to propagate uncertainty from the CelFiE decomposition. Nonetheless, it is extremely clear that the CelFiE proportions discriminating non-pregnancy from pregnancy are overwhelmingly driven by placenta

### 3.5 Application to real ALS data

Lastly, we examined cfDNA in ALS patients and age matched controls (Section 7.2 and 7.3). ALS patients represented a range of disease severities and onset sites. We first examine the overall abundance of cfDNA in cases (n=28, mean 485.9 ± 205.9pg/ul) and controls (n=25, mean 222.5 ± 58.6pg/ul) and observe a significant excess in cases (Figure 6A, *p* = 3.00 × 10^−3^). It is currently unknown what tissue or tissues are responsible for this increase. To explore possible over-represented tissues in ALS cfDNA, we apply CelFiE to 16 cases and controls for which WGBS was performed. As with the pregnancy cfDNA, we subset the WGBS data to TIM sites and then sum +/- 250 bp around the TIMs. We decompose all mixtures using the same reference tissues as Section 3.4, and one unknown (Figure 6B). Again, we choose the random restart out of 10 with the maximum log likelihood.

**Fig. 6:**
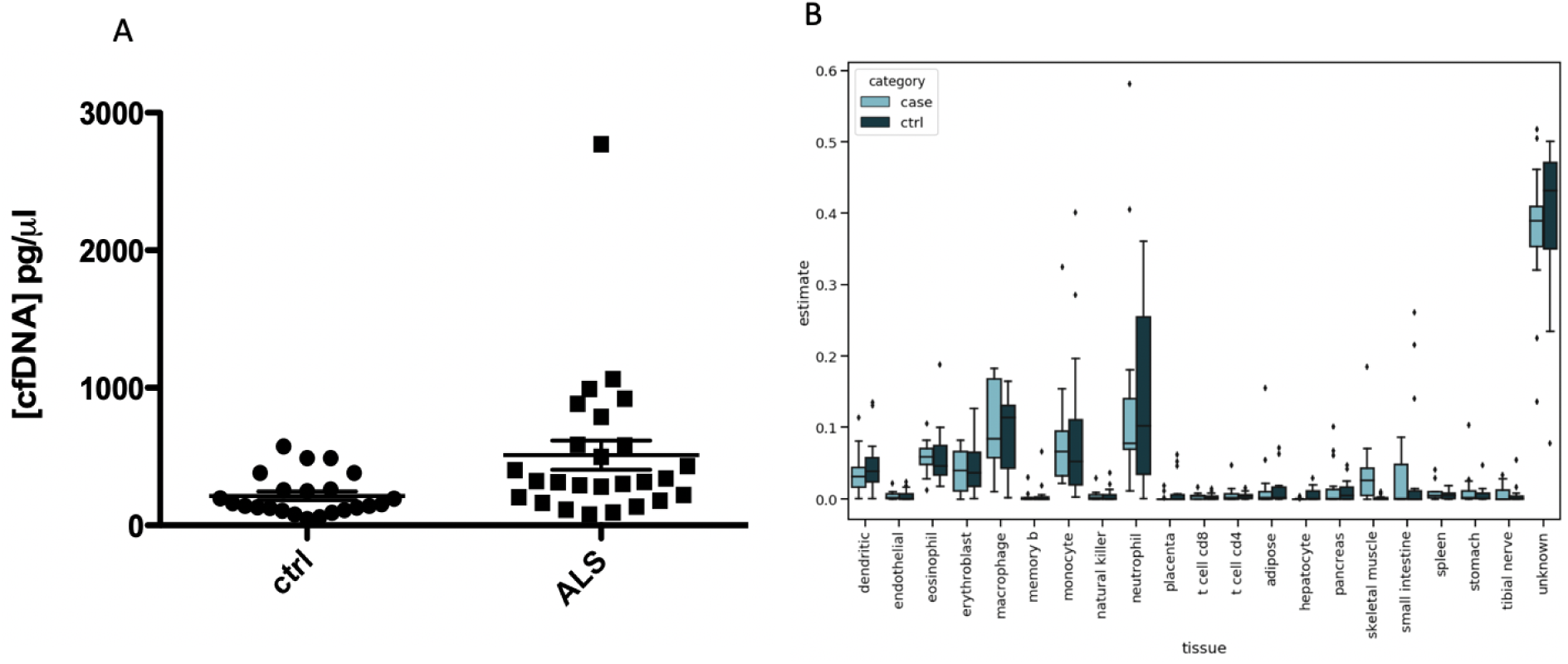
(A) CfDNA concentrations for 28 cases and 25 controls (B) CelFiE decomposition estimates for 16 ALS patients (light blue) and 16 controls (dark blue).

As expected, we find that the mixture is composed largely of white blood cells. In addition, CelFiE estimates a large unknown component, with a mean proportion of 0.38 for ALS cases and 0.39 for control samples. We next investigated specific cell type differences between cases and controls. We restrict statistical tests to three biologically relevant tissues for ALS: skeletal muscle, adipose, and tibial nerve, producing a Bonferroni threshold of 0.0167. The same grouped t-test caveats apply as in Section 3.4. Notably, we estimate a significantly higher skeletal muscle component in ALS cases, with a mean muscle proportion of 0.037 (grouped t-test *β* = 7.48, *s.e.* = 2.25, *p* = 2.67 × 10^−3^). This is more than ten times the mean proportion in controls, 0.0020. Muscle atrophy and death is a hallmark of ALS, thus, this result demonstrates CelFiE’s ability to truly learn and quantify disease relevant biology.

## 4 Discussion

During disease or increased cell turnover, elevated levels of cfDNA can be detected in the blood. For example, increases in the amount of cfDNA have been detected in patients with multiple types of cancer, autoimmune diseases, as well as acute episodes of myocardial infarction, trauma, transplantation response, and exercise [44][45][46]. Correspondingly, the utility of cfDNA as a diagnostic biomarker has been demonstrated in an increasing number of settings, including prenatal testing [47] and the detection tumor specific mutations [48][49]. Of great interest, however, is that qualitative assessments of cfDNA can now also provide information about cfDNA cellular origin [27][24][25][26]. This type of quantitative and quantitative assessment presents an individualized, unbiased approach to understanding cellular turnover over time. However, these technologies are nascent, noisy, and expensive.

In this work we presented an algorithm, CelFiE, to decompose complex cfDNA mixtures into their cell types of origin. CelFiE can accurately decompose cfDNA mixtures with low sequencing coverage in both the reference cell types and the patient cfDNA samples. Furthermore, when large cohorts are available, it can accurately estimate multiple unknown cell types, which reduces bias and increases confidence in the decomposition. Finally, the EM algorithm underlying CelFiE is computationally efficient, with iteration cost scaling linearly in the number of samples, CpG sites, and cell types.

We began by validating CelFiE extensively in simulations. Then, to demonstrate the accuracy of CelFiE on real cfDNA, we applied it to data from pregnant women. Decomposition estimates of placenta from pregnant women were significantly different from non-pregnant women. This provided a natural validation for CelFie, illustrating that it can correctly learn differences in cfDNA cell type of origin, even in real data sets.

In our study of ALS patients, we found that cfDNA levels are increased in ALS cases compared to controls. To understand what cell types are driving this difference, we applied CelFiE to the cfDNA samples, finding significantly higher skeletal muscle in patients with ALS. To the best of our knowledge, this is the first time cfDNA cell type of origin has been estimated for a neurodegenerative disease. Future work will expand on this result by expanding cohort size, and by testing for associations between cell type of origin and disease progression or severity. We may also test for associations between decomposition estimates and disease onset site. Furthermore, as cohort sizes expand, we will have power to estimate multiple unknown categories. These multiple unknown categories could be used to further subtype ALS cases. Our current results, however, are already a promising step forward, as ALS currently has no reliable biomarker. In the future, CelFiE may be used a tool for quantifying cell death in complex diseases such as ALS.

The accuracy of CelFiE depends on a number of factors including the read depth, the number and cell type-informativeness of sites considered, the abundance of key cell types, and the quantity and quality of reference data and cfDNA patient samples. Recent technologies for digesting or capturing specific regions of cfDNA [50], may allow deeper sequencing at informative CpGs. Our algorithm for selecting such TIM CpGs demonstrated marked improvement in accuracy and could be used to select sites for capture.

There are a number of areas for improvement in our approach. Many of the reference samples are complex mixtures of cell types and could be modeled as such, similar to the recent approach, FEAST [51], which modeled reference mixtures of microbial communities. Also, our WGBS simulation results showed a high degree of correlation between replicates, but we believe modeling inter-person heterogeneity will likely improve the results further in real cfDNA samples. We currently account for local correlation of CpG methylation by summing proximal CpG methylation states, but nearby CpGs may not always convey identical cell type information. Additionally, cell types are correlated in their methylation profiles and it could be interesting to consider a hierarchical model in which the composition can be considered at different levels of the cell type phylogeny. Finally, the addition of non-CpG methylation and cfDNA fragment length may provide additional sources of information about cell types of origin.

In summary, we present CelFiE, an efficient EM algorithm for decomposing cfDNA mixtures into their cell type of origin, even when the data are low count or noisy. Additionally, CelFiE can robustly estimate both known and unknown cell types in cfDNA. Overall, our work demonstrates that CelFiE could be a useful tool for quantifying cell death, applicable to biomarker discovery and disease monitoring.

## 5 Code Availability

Software available at https://github.com/christacaggiano/celfie

## 6 Acknowledgements

This work was funded by the ALS Association, ALS Finding a Cure, the ALS Biomarker Collaboration, and the UCSF Weill Award. N.Z., J.M., A.D. and C.C. are supported by NIH K25HL121295, U01HG009080, R01HG006399, R01CA227237, R03DE025665, R01ES029929, DoD W81XWH-16-2-0018, ALS Association, ALS Finding a Cure, the ALS Biomarker Collaboration, and the UCSF Weill Award.

## 7 Appendix

### 7.1 EM algorithm details for one sample

The full data likelihood is:

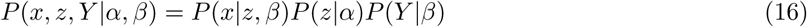

Where the first term is:

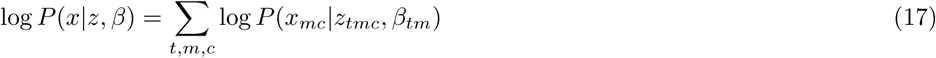

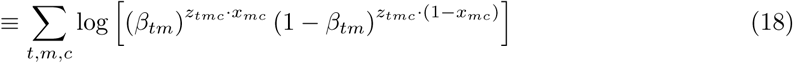

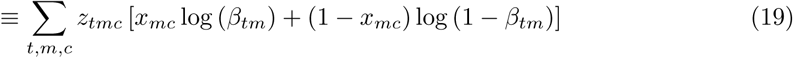

The second term is:

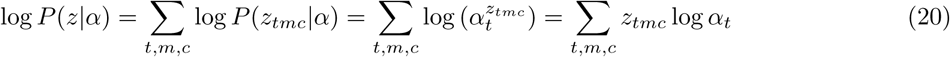

The final term is:

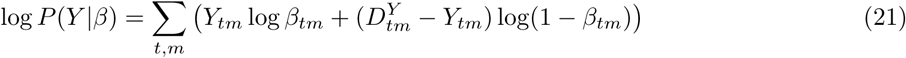

We calculate the *Q* function using the conditional distribution for *z* given some *α, β*, and the observed reads *x*:

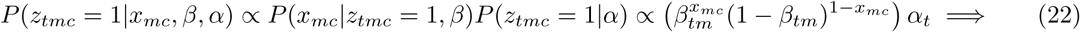

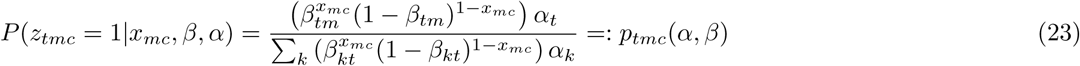

The second line follows from the fact that ∑_*t*_ *P*(*z*_*tmc*_ = 1|·) = 1, as every read must come from some cell type.

The Q-function can only have one of two values depending on the methylation state of *x*_*mc*_:

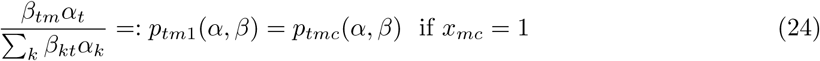

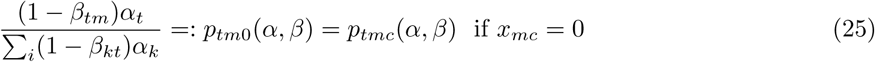

**E step:** The *Q* function is defined at iteration *i* by:

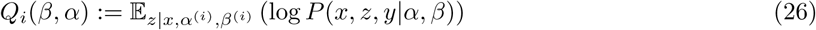

To evaluate this, we break it into three parts. Let 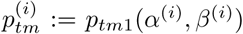–this is just the responsibility function defined above evaluated at the parameter estimates from iteration *i*. Then:

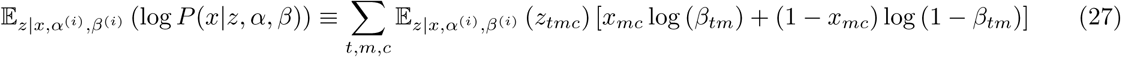

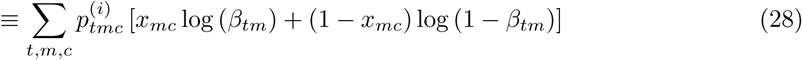

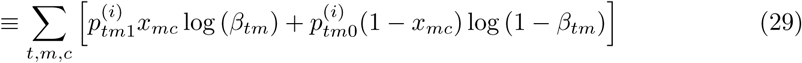

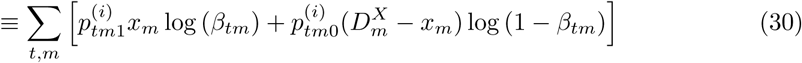

The second part is,

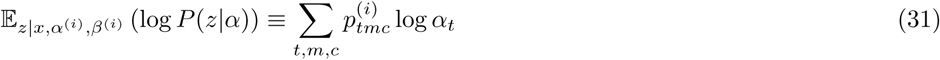

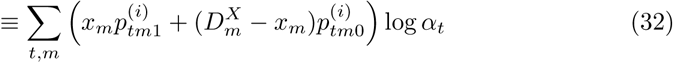

The third part is simply binomial sampling, since the cell type is known for each reference read:

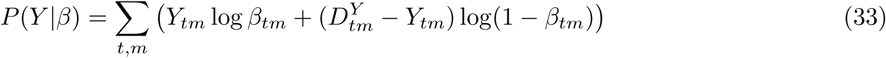

Finally, adding the three parts together:

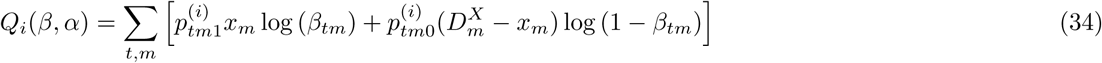

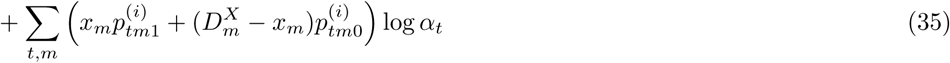

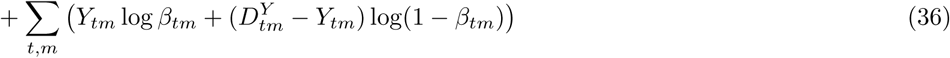

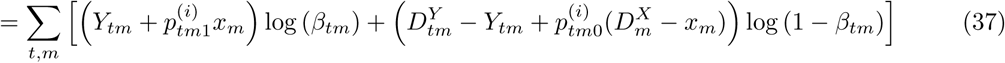

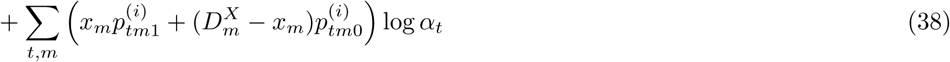

**M step:** First, let *S*_*K*_ ⊂ ℝ*K* be the probability simplex, and recall the basic fact that for any 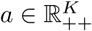 :

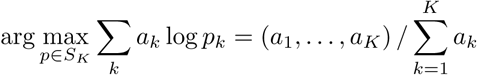

The standard way to show this is using Lagrange multipliers:

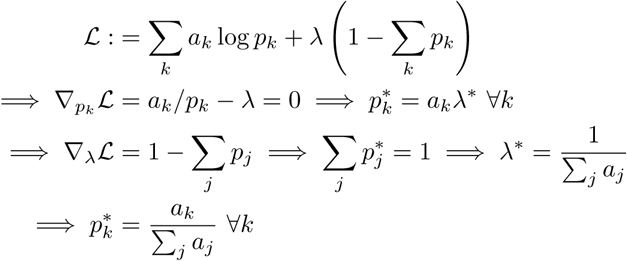

This is the only critical point of the Lagrangian, and must be a maximum since the sum of concave functions (i.e. *a*_*k*_ log *p*_*k*_) is concave; moreover, it is feasible since 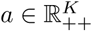 by assumption.

From these lines of basic calculus, the *α* update in (10) follows by taking 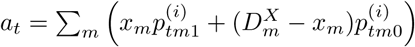. Similarly, the *β* update in (11) follows by taking 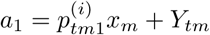 and 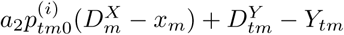.

### 7.2 Subjects

ALS patients were recruited jointly from the University of California San Francisco ALS Center and the University of Queensland ALS clinics under clinician supervision. 12 cases and 12 controls from San Francisco and 4 cases and controls from Queensland were included in this study. Controls were from non-related family members or caregivers. cfDNA was extracted after subjects were at rest for more than 30 minutes to prevent possible confounding from exercise. We collected 10 ml of whole blood from controls and 20 ml from cases, to allow for further analyses.

### 7.3 cfDNA Sequencing

Whole blood was collected in PAXgene Blood ccfDNA tubes (Qiagen, Cat. No. 768115), and centrifuged at 1,900 x g for 10 min at RT to isolate plasma. Plasma was centrifuged twice at 16,000 x g for 10 min and stored at −80*°*C until cfDNA extraction. Circulating cfDNA was extracted from 4 ml (ALS patients) or 8 ml (controls) of plasma using the QIAamp Circulating Nucleic Acid kit (Qiagen, Cat. No. 55114). cfDNA quality and concentration were assessed with an Agilent 2100 Bioanalyzer, using the Agilent High Sensitivity DNA kit (Agilent, Cat. No. 5067-4626). 10 ng of cfDNA were bisulfite-treated and purified using EZ DNA Methylation-Direct Kit (Zymo Research Cat. No D5020). Libraries for whole genome bisulfite-sequencing were generated using Accel-NGS® Methyl-Seq DNA Library Kit (Swift Biosciences, Cat. No. 30024) and Accel-NGS Methyl-Seq Dual Indexing kit (Swift Biosciences, Cat. No. 38096), with 8 cycles of indexing PCR. Libraries were quantified by qPCR with the Hyper Library Quantification kit (Kapa, Cat. No. KR0405) and paired-end sequenced on a NovaSeq 6000 System (Illumina).

### 7.4 WGBS Data Processing

Ten adult (small intestine, pancreas, monocyte, stomach, tibial nerve, macrophage, memory B cell, adipose, neutrophil and T cell) WGBS bedMethyl files were obtained from the ENCODE and BLUEPRINT project [36][38]. Each WGBS file had two biological replicates coming from distinct people. All bed file coordinates were harmonized to hg38 using hgLiftOver [52]. For each tissue or cell type, the file was restructured to report the number of methylated reads and read depth for each CpG locus. To remove sites that were non-informative of tissue of origin (i.e. the same methylation state across all tissues), CpG sites with a variance in percent methylation across tissues of less than 0.005 were removed.

### 7.5 Site Selection and Summing

#### Tissue informative markers

Only 21.8% of autosomal CpGs vary by cell type [39]. Selecting sites that do vary enriches for information on tissue of origin and reduces the EM computational burden, which scales linearly in the number of sites. We propose selecting tissue informative markers (TIMs) without curation, an approach inspired by ancestry informative markers in population genetics [53] [54].

After processing (Section 7.4) the WGBS files, one replicate per tissue was segregated into a reference matrix. This reference matrix was used to calculate TIMs. We assess whether a CpG is a TIM one locus at a time. For each CpG, the distance between the percent methylation of that cell type and the median percent methylation for that CpG was calculated. Only CpGs where the median depth was greater than 15 and had no missing data were considered. The top *N* (default=1,000) CpGs with the greatest distance per cell type were selected. TIMs provide increased accuracy in decomposition, and vastly improve computation time. We reference cell types to have overlapping TIMs (i.e., one CpG may be a TIM for both pancreas and liver). We find that TIMs perform slightly better than randomly selected sites (Figure S1). The correlation between the estimated methylation values and true WGBS methylation values for TIMs is 0.99, and the correlation for randomly selected sites is 0.90. We believe that TIMs will be especially desirable for downstream applications, where permuting random WGBS CpG sites is not feasible, or in the development of a capture panel (see Section 4).

#### Site combination

To demonstrate whether summing sites improves our ability to discriminate tissues, we create “pure” mixtures consisting of only one tissue. We select 10,000 random CpG sites per each mixture. In one experiment, we decompose the pure mixtures with no site summing. In a second, we sum +/- 250bp around the randomly selected CpG. Summing CpGs improves the performance of CelFiE (Figure S2)–the average correlation across the pure mixtures without summing is 0.79, and with summing it rises to 0.84. Overall, summing may be advantageous when data are especially low read count or noisy.

## 8 Supplemental Figures

**Fig. S1:**
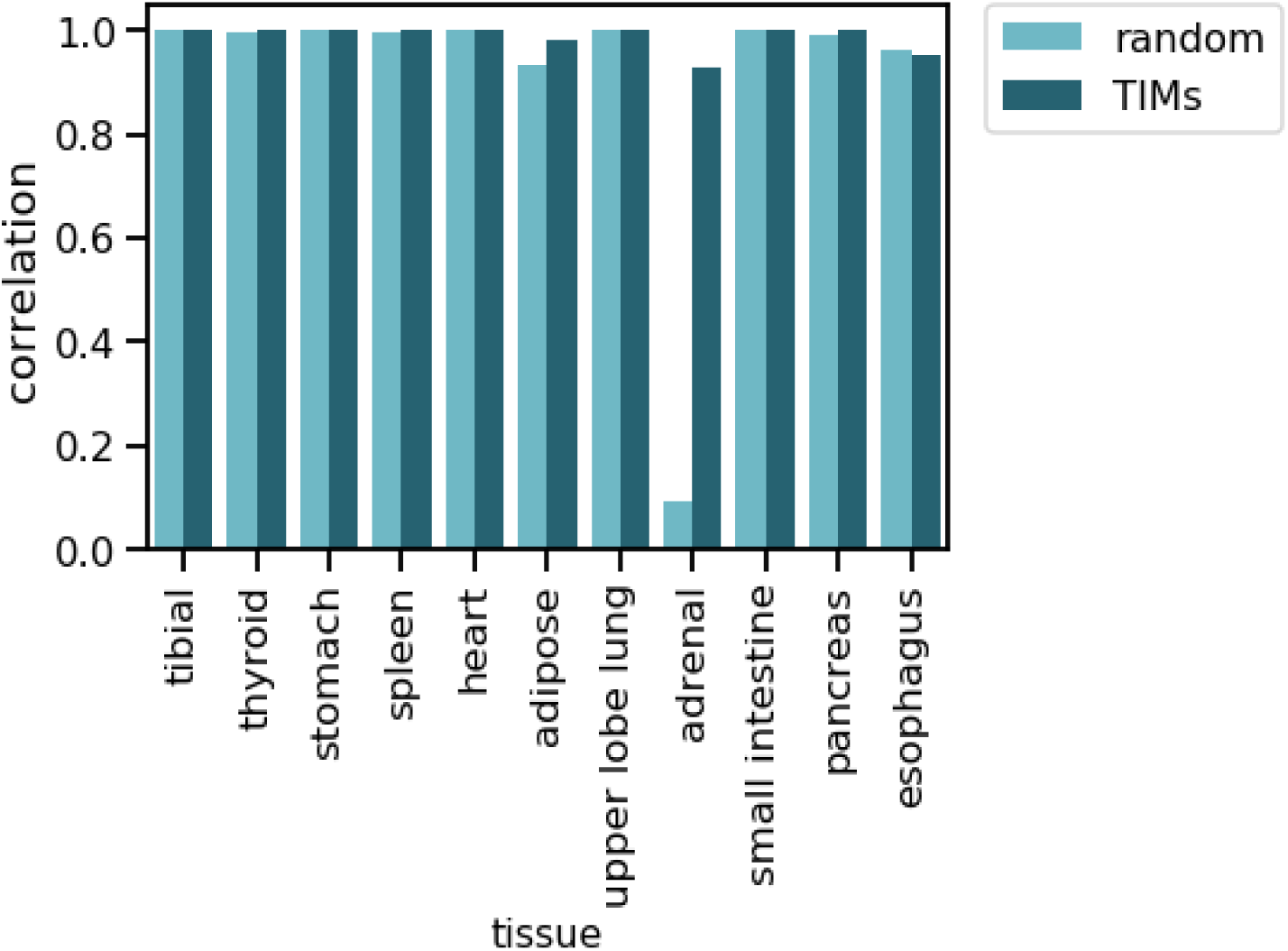
Performance of CelFie on randomly selected CpGs vs TIMs. For decomposition of mixtures 100% one tissue, we assessed the correlation between the true proportion and estimated proportion for random CpG sites (light blue), and TIMs (dark blue).

**Fig. S2:**
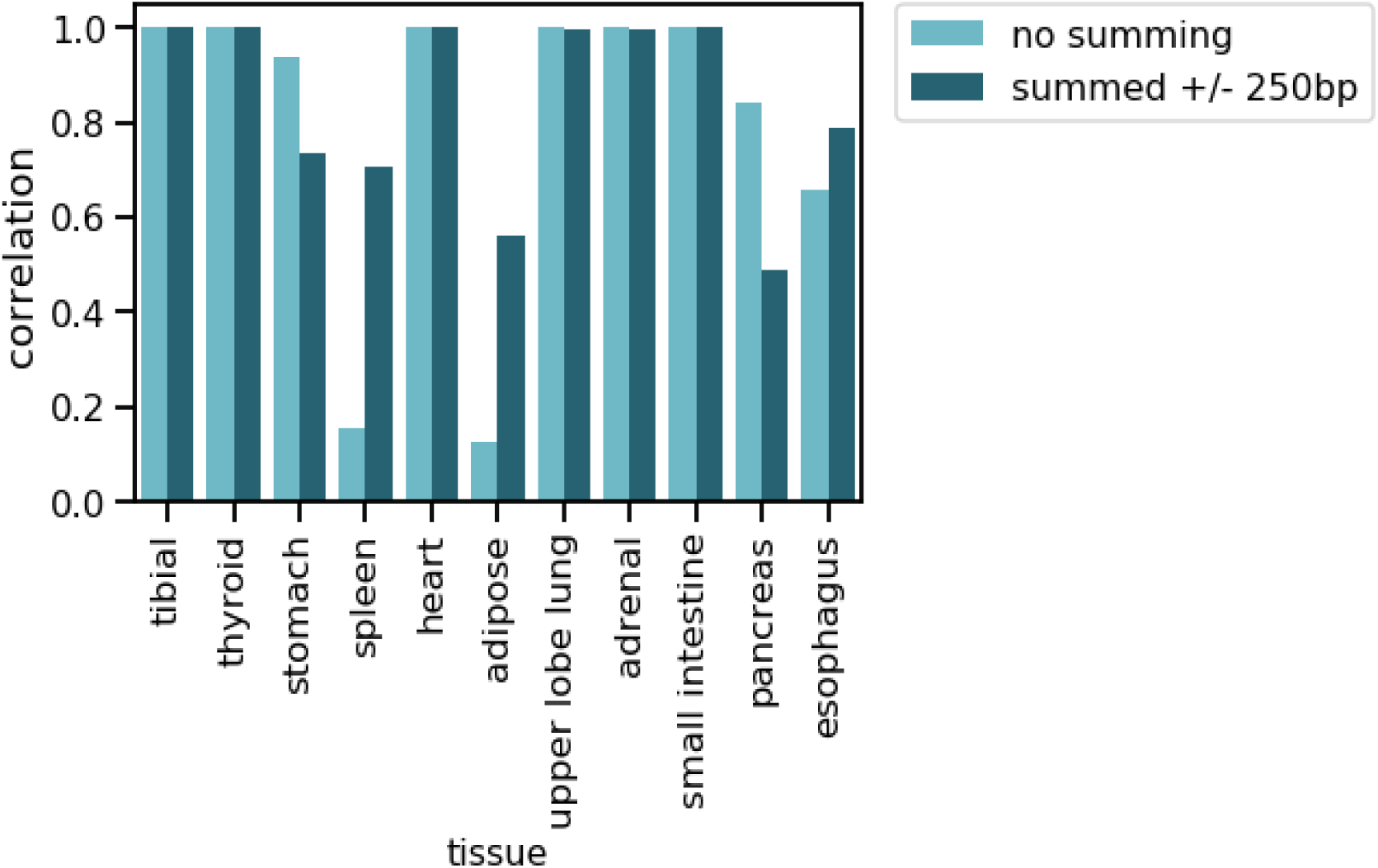
Performance of CelFie on summed versus non-summed sites. We performed decomposition of mixtures of 100% one tissue, and assessed the correlation between the true proportion and estimate for random CpG sites (light blue), and regions where reads for all CpGs +/- 250bp of that random CpG were summed (dark blue).

**Fig. S3:**
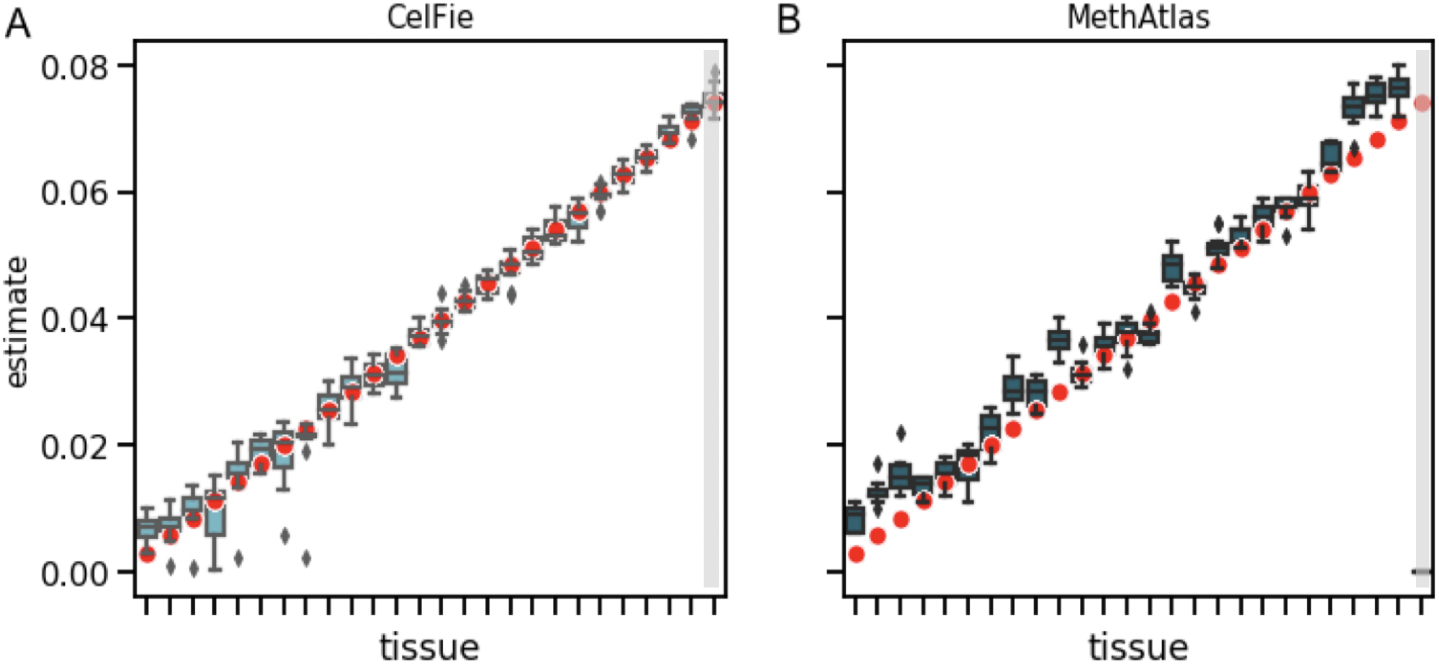
Performance of MethAtlas vs CelFiE when one cell type is missing from the reference. At 100x depth, and for 100 people, we generated cfDNA where 26 cell type are truly in the mixture. One of these cell types is then excluded from the reference (grey box). We plot the result of 50 simulation experiments for (A) CelFie and (B) MethAtlas, where the true cell type proportion is in red.

**Fig. S4:**
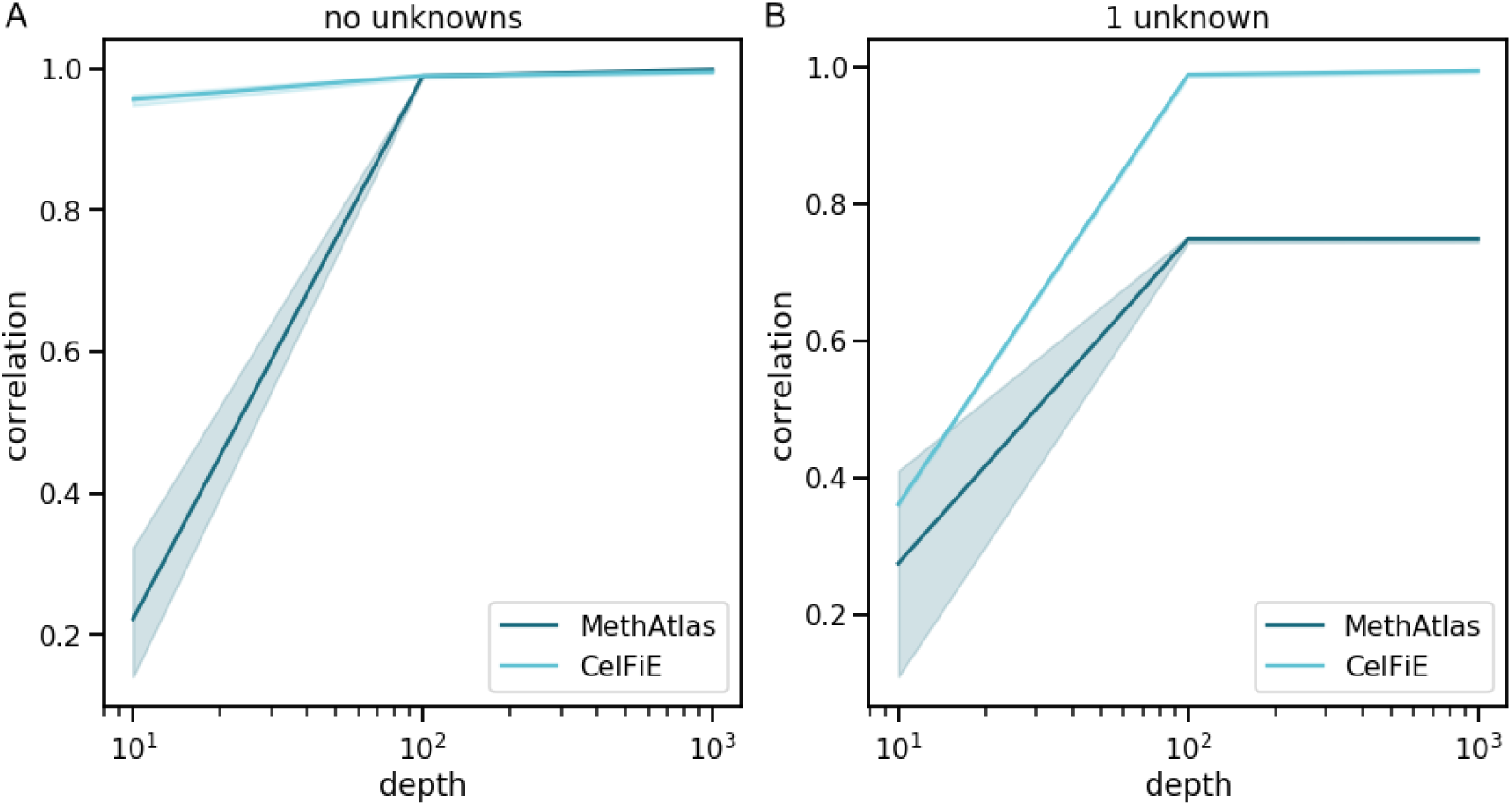
Comparison of MethAtlas and CelFiE. For a range of different depths and 100 people, we calculate the correlation between the true and estimated tissue proportion. In panel (A), all cell types are included in the reference panel. In panel (B), one cell type is missing from the reference.

**Fig. S5:**
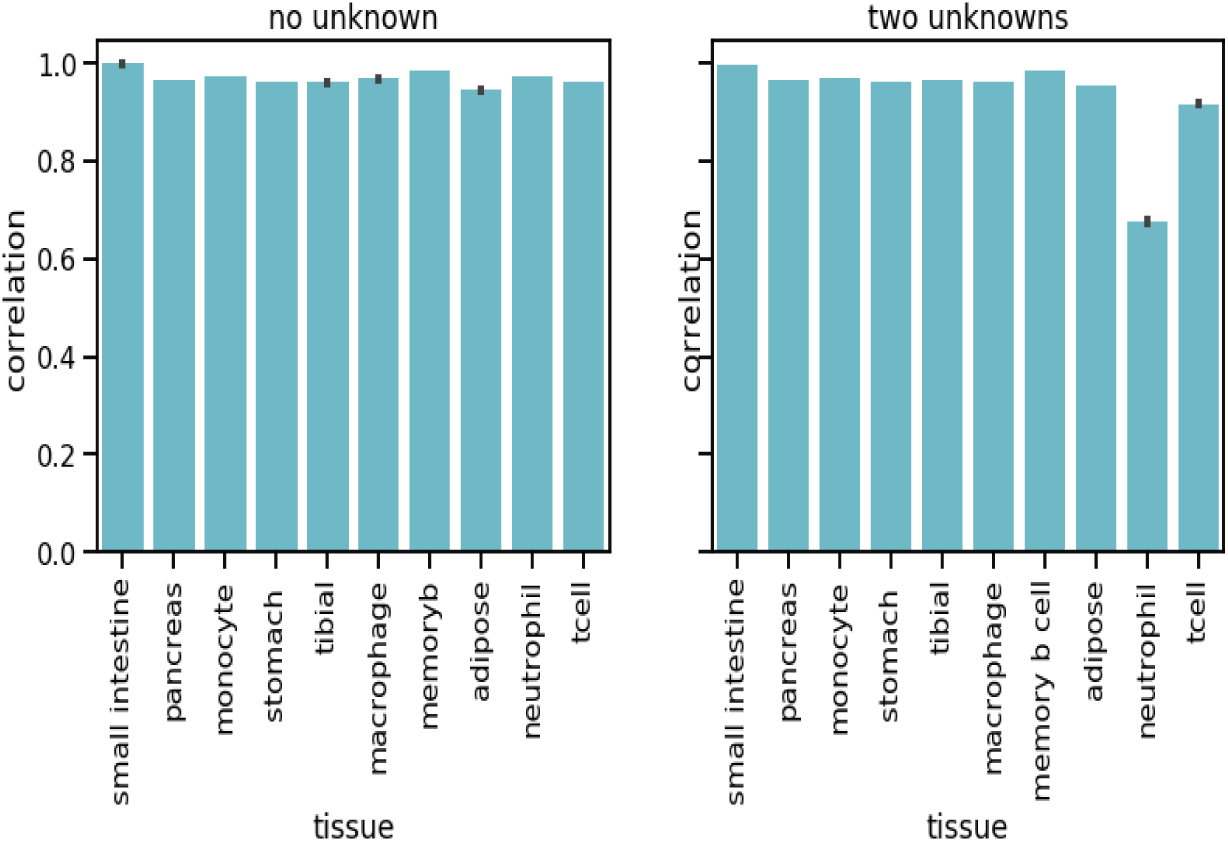
Correlation between true and estimated methylation proportion for WGBS ENCODE tissues. (A) is the correlation between the true and estimated proportions without missing tissues and (B) is with missing tissues (neutrophil and tcell).

